# Poor survival after myocardial infarction in immune deficient NXG B2m mice is alleviated by transplantation of induced pluripotent stem cell derived cardiomyocytes (iPSC-CM) formulated as spheres, but not single cells

**DOI:** 10.64898/2026.09.08.748837

**Authors:** Anne Kathrine Søgaard Terp, Ditte Gry Ellman, Sabrina Bech Mathiesen, Sara Munk Laursen, Frederik Adam Bjerre, Barbara á Grömma Ammentorp, Frederikke Gaardsvig Willadsen, Ellen Ngar Yun Poon, Charlotte Harken Jensen, Ditte Caroline Andersen

**Affiliations:** Andersen Group, Department of Clinical Biochemistry and Pharmacology, Odense University Hospital, Denmark; Department of Clinical Research, University of Southern Denmark, Odense, Denmark; Amplexa Genetics, Odense, Denmark; Hong Kong Hub of Paediatric Excellence (HK HOPE), The Chinese University of Hong Kong, Kowloon Bay, Hong Kong SAR, China; The School of Biomedical Sciences, The Chinese University of Hong Kong, Shatin, Hong Kong SAR, China; Centre for Cardiovascular Genomics and Medicine, Lui Che Woo Institute of Innovative Medicine, The Chinese University of Hong Kong, Shatin, Hong Kong SAR, China

**Keywords:** induced pluripotent stem cell-derived cardiomyocytes, iPSC-CM, NXG B2m mice, ventricular, cardiomyocytes maturation, myocardial infarction, MI, regenerative therapy

## Abstract

The human heart lacks cardiac stem cells and the ability to reestablish the loss of ventricular cardiomyocytes (vCMs) after myocardial infarction (MI) resulting in heart dysfunction. vCM replacement using intracardiac transplantation of human induced pluripotent stem cell (iPSC) derived vCMs represent a promising strategy for targeting MI, but low engraftment remains a challenge. Herein, we investigated whether formulation of ventricular iPSC-CMs (iPSC-vCM) as spheres (iPSC-vCM^Sphere^) improves cell retention and cardiac function as directly compared to single-cells (iPSC-vCM^SC^) after transplantation in a new severely immunocompromised mouse model with MI. Recipient survival immediately after MI and intervention was poor for both Vehicle (50%) and iPSC-vCM^SC^ (36%) groups, whereas 92% of iPSC-vCM^Sphere^ treated mice survived, which is comparable to non-immune deficient C57bl/6 mice (91%). All surviving and engrafted animals retained iPSC-vCMs in the infarct border zone at 8-weeks, and heart function remained similar between groups, although minor improvements were observed for iPSC-vCM^Sphere^ animals. Surprisingly, iPSC-vCM^Spheres^ were less mature than iPSC-vCM^SCs^. Thus, our data suggest that iPSC-vCM sphere formulation may offer some benefits for intracardiac delivery as compared with single cell formulated iPSC-vCMs, and thus remains a promising candidate for reestablishing the lost CMs after MI in the future.

## INTRODUCTION

According to WHO, in 2021, ischemic heart disease accounted for approximately 13% of the world’s total deaths and thus remains a leading cause of mortality worldwide.^1^ Ischemic heart disease primarily includes myocardial infarction (MI) where up to one billion cardiomyocytes (CMs) are lost.^2^ With an absence of cardiac stem cells^3^ such an extensive CM loss results in a substantial and irreversible decline in cardiac function, predisposing patients to the development of heart failure and ultimately, death.^4,5^. Currently, no curative therapy exists for advanced heart failure secondary to MI, and only the few receive a heart transplant due to the scarcity of suitable donor organs.^5^ However, regenerative medicine using induced pluripotent stem cells (iPSCs)^6^ offers new possibilities for unlimited manufacturing of iPSC-derived CMs (iPSC-CMs) to repair the MI heart upon transplantation.^7^ Already, 12 clinical trials with transplantation of iPSC-CM based products including iPSC-CM based single cell, -sheets, -spheres and patches have been registered in ClinicalTrials.gov.^8^ Yet, several challenges have emerged, and especially low cell retention, engraftment and transplantation-induced arrythmia seem major obstacles.^8–10^ Whereas ventricular arrhythmias have been linked to a fetal-like stage of the produced iPSC-CMs with immature electrophysiological properties^11^, poor cell retention and engraftment of single cell dispersed iPSC-CMs have been explained by immediate cell washout, anoikis, and immune rejection.^11,12^ Indeed, iPSC-CM based cardiac patches and sheets were successfully generated to overcome these challenges and when applied to cover the left ventricle surface in patients with heart failure, they seem to force a slight improvement in ventricular function.^8–10,13^ Yet, robust patch and sheet integration with the remaining myocardium seems limited^14^ and may indicate that these constructs remain more transient options for severe heart failure, when no other treatment is available. By contrast, single cell or sphere-formulated iPSC-CMs may offer better integration into the early remodeling MI heart, when the environment is more permissive for engraftment. Indeed, iPSC-CM spheres have been suggested to provide a 3D cellular environment in which cells maintain extensive cell-cell contacts and extracellular interactions increasing resistance to the hostile post-infarction environment, while reducing anoikis and promoting cell survival following transplantation.^11,15^ Furthermore, it is generally believed that sphere formation promotes CM maturation, electrophysiological properties, and paracrine signaling as they better mimic the 3D *in vivo* environment of the heart and hereby reduce the risk of arrythmia.^16^ Pre-clinical studies of iPSC-CM spheres in large-animal models, such as pigs and non-human primates, have shown promising results^11,17^, but direct *in vivo* evaluations after MI and particularly of the same iPSC-CMs batch formulated as single cells or spheres, remain limited^15,17^, and robust comparisons have been challenged by the low engraftment.

Herein, we thus set out to compare engraftment and heart function of iPSC-CMs formulated as single cells (iPSC-CM^SC^) or spheres (iPSC-CM^Sphere^) by transplanting them into the infarct border zone after MI. To minimize immune rejections and foster engraftment^7^, we utilized the new unreported immunodeficient NXG B2m (NOD-Prkdcscid-Il2rgtm1-B2mem1/Rj) mouse model lacking B-and T lymphocytes as well as natural killer cells. Moreover, this NXG B2m model exhibit a particularly advantageous feature with additional deficiency in β2-microglobulin, a crucial gene for the assemble of the MHC-I molecules, which in turn prevents the activation of cytotoxic T cells.^18^ Thus, in theory this model is ideally suited for human cell engraftment and direct *in vivo* comparisons of iPSC-CM^SCs^ and iPSC-CM^Spheres^.

## RESULTS

### Generation of iPSC-CMs formulated as single cells-and spheres ready for intracardiac transplantation

We used the human iPS cell line WTC-11 with a mono-allelic mEGFP tag at the c-term of the myosin light chain 2 ventricle isoform (MLC2V)^19^ for differentiation into iPSC-CMs through temporal modulation of the Wnt signaling pathway and metabolic selection^20^ followed by cryopreservation (D24) until use and subsequent maturation (Fig. 1A). At D30, high quality (Fig. S1) single-cell RNA sequencing (scRNA-seq) (Fig. 1B-F) showed that >98% (mean, SD, n=4) of the cells expressed cardiac troponin T (TNNT2; 99.3±0.3%), a-cardiac actinin (ACTC1; 99.0±1.4%), cardiac troponin C (TNNC1; 99.7±0.5%), and Titin (TTN; 98.0±1.1%) (Fig. 1C-D) reflecting high CM specification and iPSC-CM purity. In contrast, less than 0.6% of the cells contained at least one transcript of SOX2, NANOG, or POUF1 indicating that only very few if any, residual iPSCs were present in the iPSC-CM product at D30 (Fig. 1C-E). The iPSC-CM composition was highly consistent between batches (Fig. 1F) testifying to the robustness of the differentiation- and selection schedule and uniform manifold approximation and projections (UMAPs) of the distribution of CM and iPSC markers verified cluster CM identity (Fig. 1E). A large fraction (70-95%) of the cells expressed MYH7, MYL2, RYR2, and ATP5MD, demonstrating ventricular specification and CM maturity (Fig. 1C-D), and we thus refer to them as iPSC-vCMs (ventricular iPSC-CMs).

**Figure 1.**
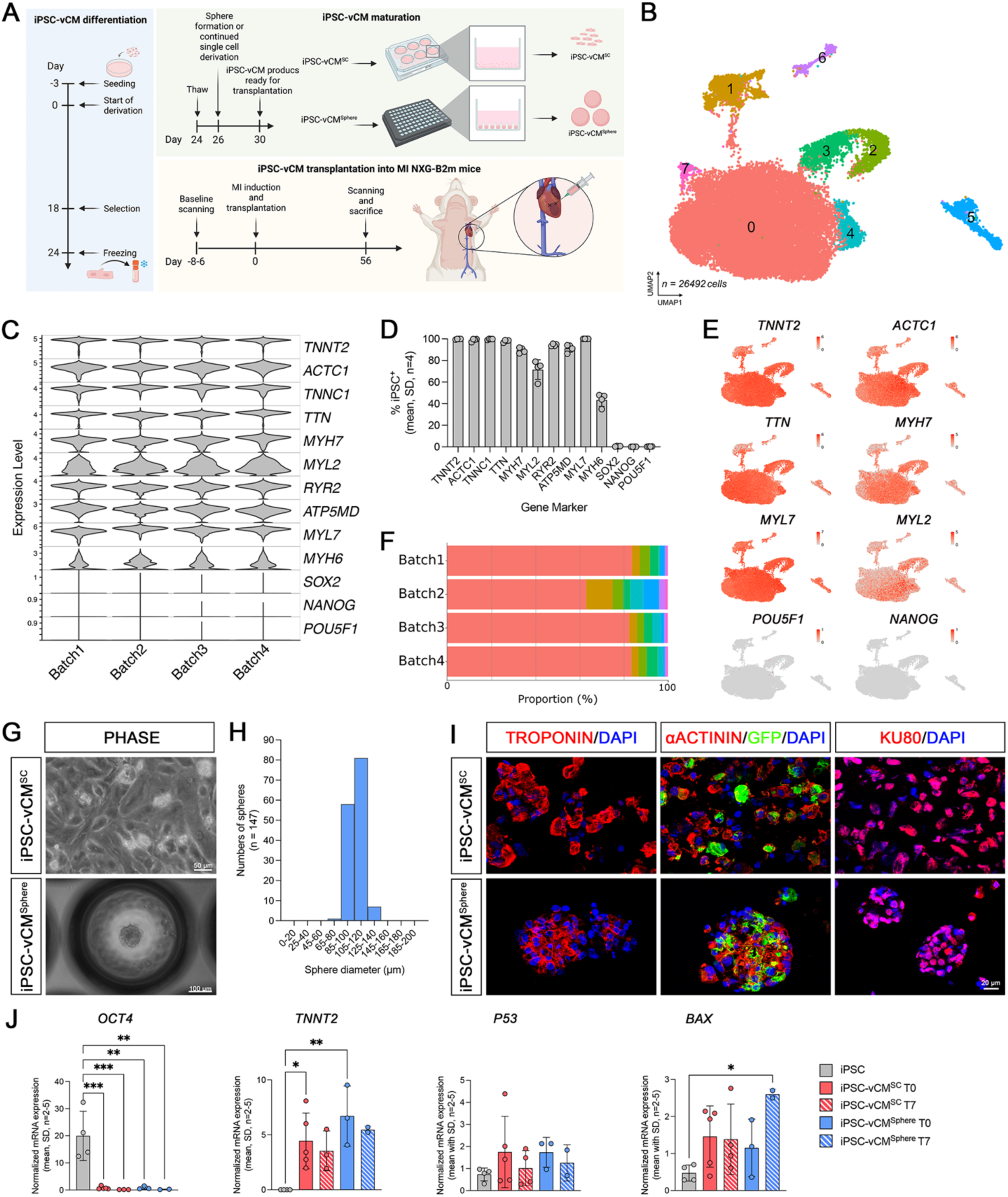
Generation and validation of single cell-(iPSC-vCM^SC^) and sphere (iPSC-vCM^Sphere^) iPSC-CM products. (A) Schematics indicating the flow in differentiation of induced pluripotent stem cells (iPSCs) to iPSC derived ventricular cardiomyocytes (iPSC-vCMs), iPSC-vCM maturation and sphere (iPSC-vCM^Sphere^) formation, and single cell (iPSC-vCM^SC^) or iPSC-vCM^Sphere^ injection into NXG-B2m mice with myocardial infarction (MI). (B) Uniform manifold approximation and projection (UMAP) plot of day (D) 30 CMs showing the eight (0-7) different CM clusters. (C) Violin-plots showing the expressions levels of TNNT2, ACTC1, TNNC1, TTN, MYH7, MYL2, RYR2, NKX2.5, ATP5MD, MYL7, MYH6, SOX2, NANOG, and POU5F1 of the four different CM batches. (D) Percentage of sequenced iPSCs positive for cardiac-(TNNT2, ACTC1, TNNC1, TTN, MYH7, MYL2, RYR2, ATP5MD, MYL7, MYH6) and pluripotency markers (SOX2, NANOG, and POU5F1), as average of the four batches. (E) UMAPs of the distribution of TNNT2, ACTA1, TTN, MYH7, MYL7, MYL2, POU5F1 and NANOG in the eight identified CM clusters. (F) Proportional (%) distribution of iPSCs in the eight CM clusters for the for batches. (G) Representative phase pictures of iPSC-vCM^SC^ and iPSC-vCM^Sphere^, scalebar 50 and 100µm, respectively. (H) Sphere diameter of representative iPSC-vCM^Spheres^ used for injection in NXG-B2m mice with MI (n=147). (I) Representative TROPONIN T/DAPI, α-ACTININ/GFP/DAPI, and KU80/DAPI stained iPSC-vCM^SC^ (upper panel) and iPSC-vCM^Sphere^ (lower panel). (J) Normalized mRNA levels of *OCT*, *TNNT2*, *P53*, and *BAX* in iPSC, iPSC-vCM^SC^ at T0 (sample from cell-suspension at the start of transplantation) and T7 (sample from cell-suspension after the last transplantation), and iPSC-vCM^Sphere^ at T0 and T7. *≤0.05: **: p<0.01, ***: p<0.001.

To compare iPSC-vCM^SC^ and iPSC-vCM^Sphere^ formulations of iPSC-vCMs for *in vivo* engraftment and efficacy after MI, D24 iPSC-vCMs were thawed and seeded, and at D26 either maintained as iPSC-vCM^SC^ to facilitate further maturation or aggregated into multicellular iPSC-vCM^Spheres^ using specialized microcavity plates during maturation (Fig. 1A, G). To enable later robust iPSC-vCM^Sphere^ intracardiac delivery through a 29-gauge needle, we found spheroids with a mean diameter of 104.7µm±10.2 (SD, n=147) and an average of approximately 250 iPSC-vCMs per spheroid to be optimal and easily reproducible regarding shape and size (Fig. 1H and Fig. S2). Immunofluorescence analysis of cryo-embedded iPSC-vCM^SC^ and iPSC-vCM^Sphere^ showed expression of TROPONIN T and α-ACTININ as well as GFP/MYL2 confirming CM specification with a ventricular fate (Fig. 1I). Distinctive KU80 signals substantiated the human cell origin important for *in vivo* tracking after transplantation into mice (Fig. 1I). For transplantation, cells were maintained at room temperature in a vehicle survival cocktail for up to 6 h, with no significant differences between groups and time in levels of P53 or BAX, key regulators of apoptosis (Fig. 1J). Collectively, these findings demonstrated the successful generation of highly pure iPSC-vCM^SC^ and iPSC-vCM^Sphere^ formulated cell products suitable for intracardiac transplantation after MI.

### NXG B2m mice are exceptionally fragile after MI, but survival increases with iPSC-vCM^Sphere^ treatment

A total of 49 female NXG B2m mice were included in the study at baseline (week -1) for functional heart scanning (Fig. 1A and 2A). Mice surviving initial scanning and intubation/anesthesia underwent MI and were randomly assigned to: Vehicle (PBS^Cocktail^) (n=12,) iPSC-vCM^SC^ (n=14), iPSC-vCM^Sphere^ (n=12) for transplantations. A sham group (n=2) was used for surgery assessment. Mice from each group were mixed and distributed equally on all experimental days. However, very quickly, we realized that the NXG B2m mice were extremely fragile upon MI and/or transplantation, and many animals died and were excluded from the study (Fig. 2A). We were unable to include more NXG B2m animals to compensate for this loss due to ethical restrictions. Furthermore, as a precaution, the planned week 1 functional heart scanning was omitted due to death of the first mouse scanned (Fig. 2A). At termination, Kaplan-Meier survival curves clearly reflected the observed abrupt death upon intervention in the Vehicle group and further worsened in the iPSC-vCM^SC^ group, whereas survival was surprisingly and significantly (p=0.0005) higher in the iPSC-vCM^Spheres^ group and not significantly different from sham mice (Fig. 2B). Indeed, NXG B2m recipient survival within the first 2 days after MI and intervention was only 50- and 36% for Vehicle and iPSC-vCM^SC^ groups, respectively, whereas 92% of iPSC-vCM^Sphere^ treated NXG B2m mice survived. The latter was comparable to immune-compatible C57bl/6 mice after MI and injections of either Vehicle (PBS) or iPSC-vCMs showing survival rates >90% (Fig. 2C). Comprehensive analysis of animals dying before study termination and sectioning throughout the NXG B2m hearts with tracking of human iPSC-vCMs via KU80 (Fig. 1H), showed no difference in the locations of transplanted iPSC-vCM^SCs^ and iPSC-vCM^Spheres^ (Fig. 2D). Only one mouse showed iPSC-vCM^SCs^ inside the ventricle lumen (Fig. 2D). This excluded the possibility that iPSC-vCM^SC^ NXG B2m mice mainly died from embolism. Microscopic examination and stereological analyses of animals included at study termination (8 weeks) showed that all NXG B2m hearts independent of intervention, exhibited a very thin, almost transparent wall of the left ventricle, with blood inside the ventricle lumen being visible (Fig. 2E). This is very different from MI hearts in normal mice like C57bl/6, where extensive remodeling with fibrosis after MI facilitates closure of the wound and prevents excessive wall thinning.^21^ To assess this general vulnerability of the NXG B2m to MI, we compared echocardiography scanning from Vehicle treated-C57bl/6 and NXG B2m mice at baseline and at follow up after MI. No differences were present functionally or in wall thickness between mouse strains at baseline, but NXG B2m mice were substantially compromised in all echocardiography parameters at follow up (Fig. 2F and Fig. S3). This underscores that the NXG B2m strain by itself is extremely fragile and vulnerable after MI independent of iPSC-vCM interventions.

**Figure 2.**
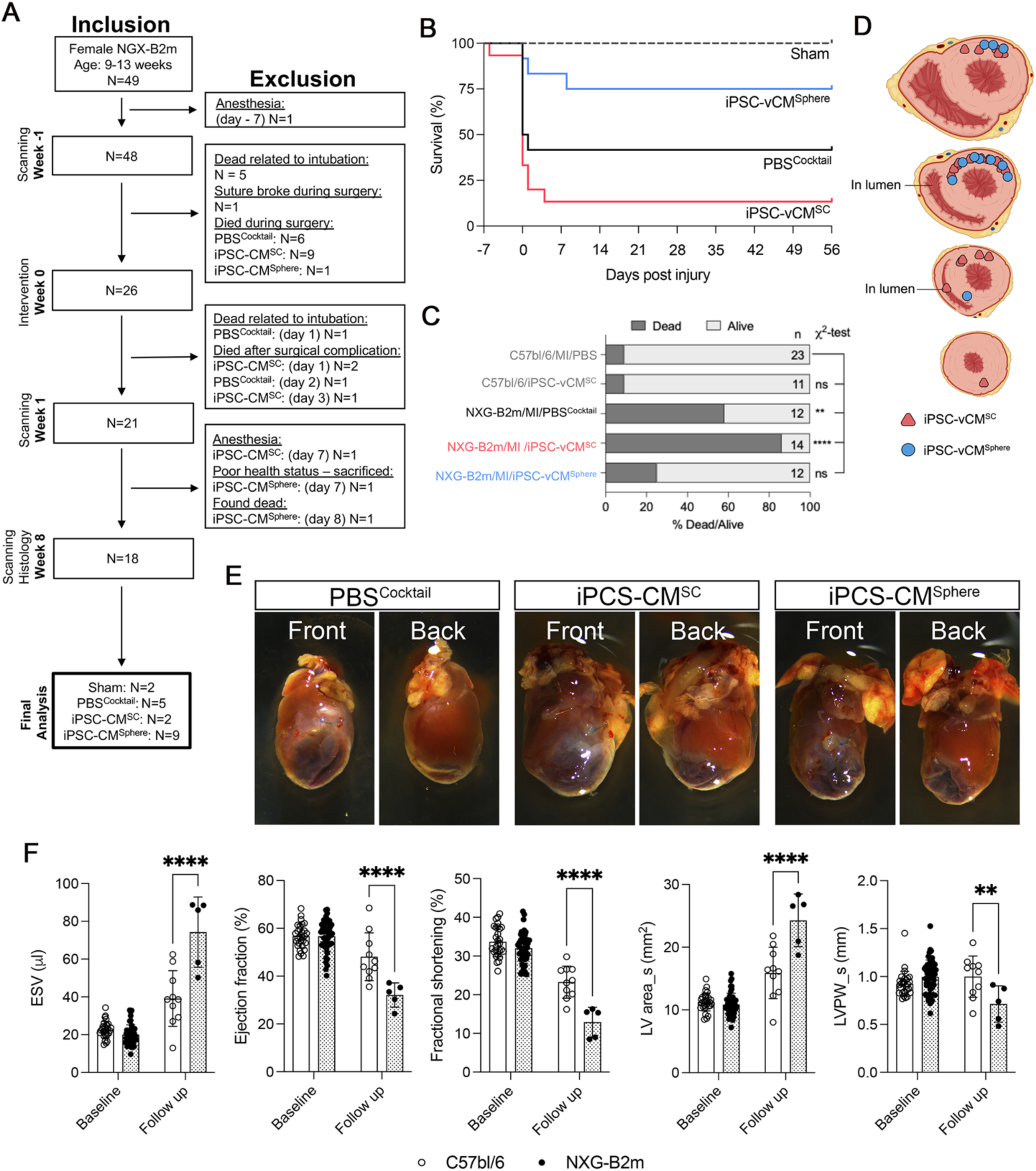
*In vivo* study design, survival, and functional performance of NXG-B2m mice after MI and iPSC-vCM intervention. (A) Overview of *in vivo* timeline, and inclusion and exclusion of NXG-B2m mice after MI and cell transplantation. (B) Kaplan-Meier survival curve showing the survival rate of Sham, PBS^Cocktail^, iPSC-vCM^SC^, and iPSC-vCM^Sphere^ groups. (C) Survival rate of C57bl/6 mice after MI and PBS injection, C57bl/6 mice after transplantation of iPSC-vCM^SC^, NXG B2m mice after MI and injection of PBS^Cocktail^, NXG-b2m mice after MI and transplantation of iPSC-vCM^SC^, and NXG B2m mice after MI and transplantation of iPSC-vCM^Sphere^. (D) Schematic overview of detected iPSC-vCM^SC^ (red triangles) and iPSC-vCM^Sphere^ (blue dots) in NXG B2m mice dying before study termination. (E) Representative pictures of the front and the back of NXG B2m mouse hearts transplanted with PBS^Cocktail^, iPSC-vCM^SC^, or iPSC-vCM^Sphere^, respectively, at termination 8 weeks after MI and intervention. (F) Functional comparison of end systolic volume (ESV), ejection fraction, fractional shortening, left ventricle (LV) area, and left ventricle posterior wall (LVPW) thickness in C57bl/6 and NXG B2m mice at baseline and follow-up after MI and vehicle injection. **: p<0.01, ***: p<0.001, ****: p<0.0001.

### iPSC-vCM^SC^ and iPSC-vCM^Sphere^ engraft in the MI border zone, with no overall difference in functional output or infarct size despite a difference in maturity

Eight weeks after MI and intervention, we detected iPSC-vCMs in all surviving iPSC-vCM^SC^ and iPSC-vCM^Sphere^ NXG B2m mice as tracked by visualization of human nuclei (Fig. 3A-D). Regardless of an uneven number of included mice for analysis, we found no apparent difference in the localization of iPSC-vCM^SC^ and iPSC-vCM^Sphere^ in the hearts (Fig. 3A-D). Several engrafted iPSC-vCMs resided in the border zone of the MI (Fig. 3B-C) in the anterior and posterior wall of the left ventricle. Both iPSC-vCM^SC^ and iPSC-vCM^Sphere^ were retained in relative proximity to the injection site, with no appearances in the apex, but still distributed across many levels between the apex and base of the heart (Fig. 3D). Importantly, we did not observe any biodistribution of neither iPSC-vCM^SC^ nor iPSC-vCM^Sphere^ in kidney, liver, lung and spleen (Fig. 3E) using a highly sensitive Alu Y sequence assay validated by spike-in (Fig. S4). Although some minor indications of better functional improvements for iPSC-vCM^Sphere^ treated mice were observed, heart function (Fig. 3F) and infarct size (Fig. 3G) were equally compromised over time after MI and not different from Vehicle injected animals.

**Figure 3.**
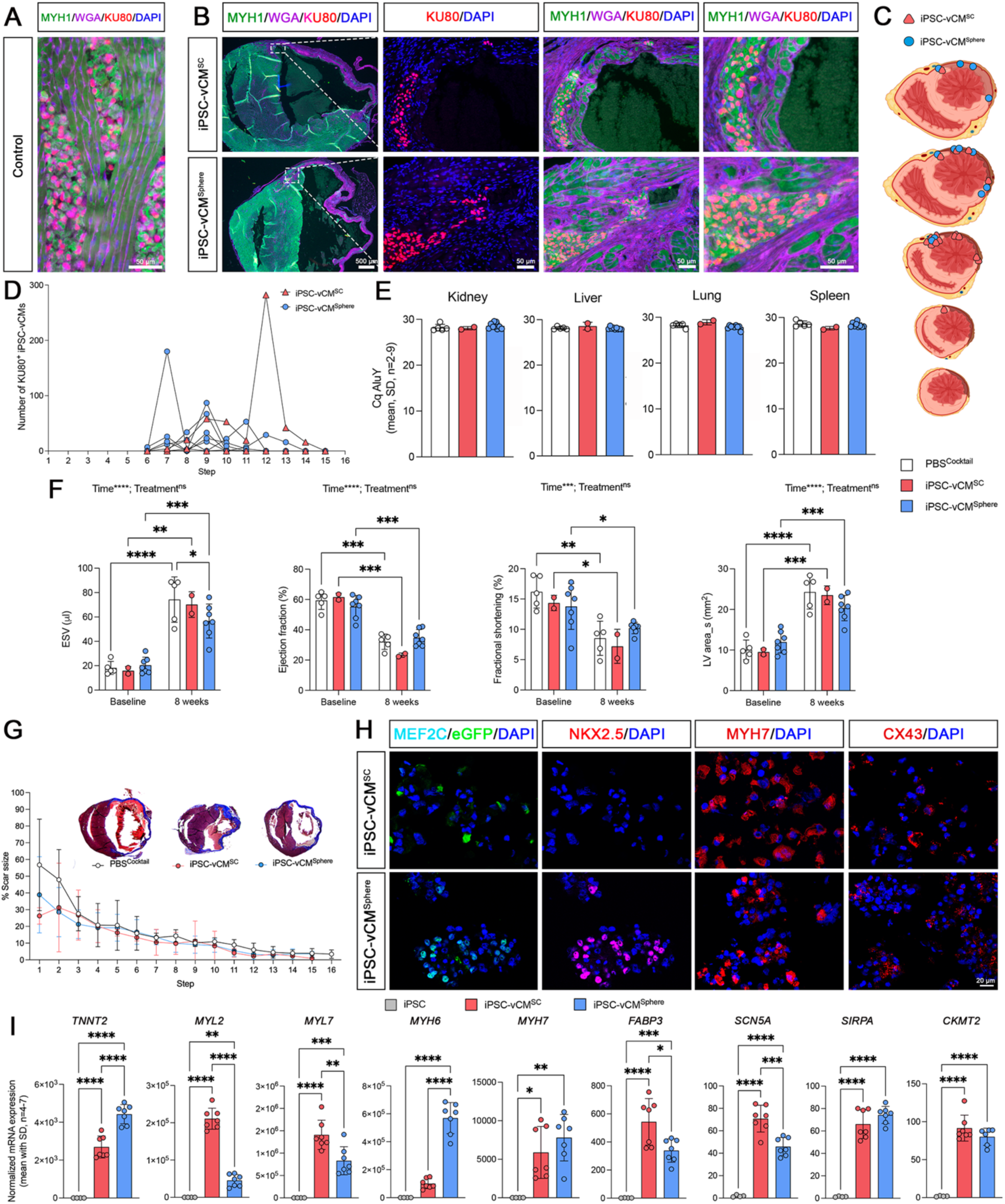
iPSC-vCM^SC^ and iPSC-vCM^Sphere^ engraftment, biodistribution, functional recovery, scar formation in NXG B2m mice and maturity characterization of injected iPSC-vCM^SC^ and iPSC-vCM^Sphere^. (A) Positive control mouse heart with engrafted induced pluripotent stem cell derived ventricular cardiomyocytes (iPSC-vCMs) (MYH1: green; Wheat germ agglutinin (WGA): purple; KU80: red; DAPI: blue). Scalebar: 50µm (B) Representative pictures of NXG B2m hearts transplanted with iPSC-vCM^SC^ or iPSC-vCM^Sphere^ after 8 weeks (MYH1: green; Wheat germ agglutinin (WGA): purple; KU80: red; DAPI: blue). Scalebars: 500µm (left panel), 50µm (middle and right panels). (C) Schematic overview of the presence of transplanted iPSC-vCM^SC^ (red triangles) and iPSC-vCM^Sphere^ (blue dots) after 8 weeks. (D) Presence of transplanted iPSC-vCM^SC^/iPSC-vCM^Sphere^ in the different steps in sectioned NXG B2m hearts at 8 weeks. (E) Cq AluY values in genomic DNA from kidney, liver, lung, and spleen from NXG B2m mice transplanted with either PBS^Cocktail^, iPSC-vCM^SC^, or iPSC-vCM^Sphere^, after 8 weeks. (F) Heart function (end systolic volume (ESV), ejection fraction, fractional shortening, left ventricular (LV) area in systole, and left ventricular posterior wall (LVPW) thickness in systole) of NXG B2m mice heart transplanted with either PBS^Cocktail^, iPSC-vCM^SC^, or iPSC-vCM^Sphere^ at baseline and after 8 weeks. (G) Stepwise scar size in NXG B2m mice heart injected with either PBS^Cocktail^, iPSC-vCM^SC^, or iPSC-vCM^Sphere^ after 8 weeks. (H) Representative pictures of injected iPSC-vCM^SC^ (upper panel) or iPSC-vCM^Sphere^ (lower panel) (MEF2C: cyan; eGFP: green; NKX2.5, Myh7, CX43: red; DAPI: blue) (I) Normalize mRNA expression of TNNT2, MYL2, MYL7, MYH6, MYH7, FABP3, SCN5A, SIRPA, and CKMT2 in iPSC, iPSC-vCM^SC^, and iPSC-CM^Sphere^. *≤0.05: **: p<0.01, ***: p<0.001, ****: p<0.0001.

Thus, since overall engraftment and efficacy were similar between intervention groups, we speculated how to explain the differences in survival and cardiac arrest after transplantation (Fig. 2A-C). From the Vehicle data comparing NXG B2m mice with normal mice (Fig. 2F), it seems reasonable to conclude that MI by itself has a major impact on heart remodeling and function in the NXG B2m mice as also known from similar studies. But it is also possible that fluid injection in itself may affect the stability of the NXG B2m heart specifically, triggering mechanical induced arrhythmia.^22^ However, how iPSC-vCM^Sphere^ treatment counteracted the poor survival of NXG B2m mice seemed highly enigmatic. Since transplantation-induced arrhythmia of iPSC-vCMs is a challenge normally associated with low maturity level of the injected iPSC-vCMs^11^, we used qPCR to compare expression levels of maturity markers between the two cell formulations. Interestingly, we observed that iPSC-vCM^SCs^ expressed higher levels of *MYL2, MYL7, FABP3, SCN5A*, all genes linked to maturity, whereas expressions of the cardiac progenitor- and immaturity markers *MYH6*, NKX2.5 and MEF2C were more pronounced in the iPSC-vCM^Spheres^ (Fig. 3H-I). Both iPSC-vCM^SCs^ and iPSC-vCM^Spheres^ expressed MYH7 and CX43 reflecting a capacity for synchronized cardiac contraction. These data surprisingly showed that the iPSC-vCM^Sphere^ product was less mature than the iPSC-vCM^SC^ formulation, which may affect the risk of arrhythmia induction and sudden death in the NXG B2m mice after MI.

## DISCUSSION

Herein, we found that poor survival of severely immunodeficient NXG B2m mice after MI was rescued by intracardiac transplantation of iPSC-vCMs formulated as spheres, but not single cells. In general, survival after MI is affected by many parameters. Our survival rate of 90% in normal healthy mice after MI is accepted as high. Thus, we do not as such consider technical issues to underlie the high risk of sudden cardiac arrest we observed in NXG B2m mice after MI for both Vehicle and iPSC-vCM^SC^ treated animals. A mouse age below 12 weeks has previously been linked to lower survival after MI in the NOD-SCID mouse that also lacks B cells, T cells and natural killer cells.^23^ Our mice had a mean age of 13±1 weeks at the time of MI, but whether the additional absence of MHC-I in the NXG B2m mouse further increases the risk of sudden cardiac arrest is unknown but seems likely. It is however evident that the immune system is essential for extracellular matrix decomposition and scar formation^24^ and immune deficient mice may thus risk wall thinning after MI. This is clearly supported by our data where immune deficient mice exhibited cardiac properties similar to normal mice at baseline, but obtained a severe phenotype after MI, where heart dysfunction was exaggerated. In agreement, we found that the left ventricular wall was poorly remodeled, and wall thinning was obvious 8 weeks after MI for all groups. Since all mice, except for shams, received injections with fluid±iPSC-CMs, we cannot rule out that the high sudden cardiac arrest at least partly could be a consequence of MI combined with fluid injection where NXG B2m mice may be more susceptible to fluid stress. Yet, this is contradicted by the observation that iPSC-vCM^Sphere^ transplantations rescued the NXG B2m mice from sudden cardiac arrest achieving survival rates corresponding to that of normal mice after MI. iPSC-vCM^Sphere^ products have previously been shown to mitigate early loss from the injection site due to leakage, myocardial contraction and washout into the circulation supposably by maintaining cells as cohesive aggregates, resulting in improved early retention.^25^ Only two studies have compared *in vivo* safety and efficacy for iPSC-vCM^SCs^ and iPSC-vCM^Spheres^, but low engraftment was evident in both studies despite using immune deficient animals or immune suppressive therapy^15,17^. Yet, the study by Park et al. did report that their survival rates corresponded to 55%, 55% and 91% for vehicle (PBS), iPSC-vCM^SCs^ and iPSC-vCM^Spheres^, respectively^15^. These data are very similar to our survival data, and together our two studies underscore that sphere formulation seem to benefit iPSC-vCM delivery and recipient survival. Despite some minor indications that cardiac function was less compromised in iPSC-vCM^Sphere^ treated NXG B2m mice, the low number of surviving Vehicle and iPSC-vCM^SC^ treated animals prevents any robust conclusions on efficacy at this stage. Unfortunately, including additional NXG B2m mice into the study was not an option due to ethical restrictions with the observed low survival. Future comparisons of iPSC-vCM^SC^ and iPSC-vCM^Sphere^ products in regard to efficacy after engraftment in the MI heart may alternatively be performed in less immune deficient mice with hypoimmunogenic iPSCs.^26^ Yet, we did observe that iPSC-vCM^Spheres^ are less mature than iPSC-vCM^SC^ which is surprising since several studies have suggested that sphere formation increases maturity^16^ which is important to avoid transplantation-induced arrhythmia.^7,11,17^ Our results directly add to this controversy indicating that iPSC-vCM maturity not necessarily promotes engraftment and recipient survival. It is likely, that the observed immaturity of the iPSC-vCM^Sphere^ product results in a less beating CM phenotype that lowers the risk of a beating mismatch upon transplantation and hereby the risk of sudden cardiac arrest from arrhythmia. More studies are however required to explore this in detail.

In agreement with the theory, the severe immune deficient environment explored herein substantiated iPSC-vCM engraftment as compared to our transplantations in normal mice with immune suppressive therapy, where iPSC-vCM cannot be detected already after 1-2 weeks in agreement with others.^27,28^ The clear border zone localization of the engrafted iPSC-vCMs independent of formulation also nicely demonstrates that transplanted iPSC-vCMs seldomly survive the hostile necrotic area inside the infarct even in immune deficient mice. In a clinical perspective this underlines that iPSC-vCM transplantations should be timed carefully to avoid the necrotic response occurring in the first days after MI.^29^ Timing iPSC-vCM transplantations after necrosis, but before fibrosis may enable better engraftment directly in the infarct zone, which is important as iPSC-vCMs is considered non-migratory.^30^

Our study clearly exhibits some limitations caused by the low survival of recipients, but this in itself provides some invaluable information to the field on the use of new NXG B2m mouse and similar severely immune deficient mouse strains for assessing cell transplantations upon tissue injury. Moreover, we used only one iPS cell line and therefore cannot generalize our conclusions but notice that the WTC11-MLC2v-GFP is commonly used for heart research.^31,32^

In summary, our data suggest that iPSC-vCM sphere formulation may offer some benefits for intracardiac delivery as compared with single cell formulated iPSC-vCMs and thus remains a promising candidate for reestablishing the lost CMs after MI in the future.

## MATERIALS & METHODS

### Animals

Fortynine female NXG B2m (NOD-Prkdc^scid^-Il2rg^tm^^1^-B2m^em^^1^/Rj) mice were purchased from Janvier (Le Genest-Saint-Isle, Saint Berthevin Cedex, France) and housed in a specific pathogen-environment at the Biomedical Laboratory (University of Southern Denmark, Odense, Denmark) in groups of 4 mice in individually ventilated cages in a 12/12 hour light/dark cycle with free access to food and water. At the time of baseline scanning the mice were between 9 and 13 weeks old. All animal experiments were approved by the Danish Counsil for Supervision with Experimental Animals (# 2022-15-0201-01119).

### Derivation of iPSC-vCM

WTC11-MLC2v-GFP (Allen Institute for Cell Science available from Coriell Institute for Medical Research, AICS-0060-027) iPSC were cultured in mTeSR medium (Stem Cell Technologies, 100-0274) on growth factor reduced Matrigel (Corning, 356231) and routinely passaged using Accutase (Thermo Fisher, A11105-01) with mTeSR+ supplemented with Y27632 2HCl (Selleckchem, S1049). CM differentiation was performed using a small molecule-based differentiation protocol based on temporal modulation of Wnt signaling and metabolic selection as previously described^20^. On differentiation D26 iPSC-vCMs were trypsinized and plated in Corning® Elplasia Plates (VWR CORN4441) to generate iPSC-vCM^Spheres^. Cells were harvested for characterization or transplantation at D30.

### ScRNA-seq

ScRNAseq was performed as previously described^33^. Briefly, single-cell libraries were generated using the Chromium Next GEM Single Cell 3’ GEM Kit v3 (10X Genomics, PN-1000123), following the manufactures instructions and targeting 10,000 loaded cells per run. cDNA libraries were sequenced using the Illumina NovaSeq 6000 (Illumina inc.). Data analysis was performed as previously described,^34^ with minor cell-type-specific modifications. In short, sequenced iPSC-CM data was processed using CellRanger (9.0.1) with default parameters using the GRCh38 (GENCODE v32/Ensembl98) reference genome. Downstream analysis was performed in R (v4.5.2) with use of the Seurat package (v5).^35^ Genes expressed in fewer than 3 cells were removed, as were cells falling outside IQR × 1.5 bounds (minimum 300 genes). iPSC-CM batches were normalized individually using (LogNormalize, scale factor 10,000), and the top 2000 most variable genes were used for dimensional reduction. Batch effects were corrected using Harmony (default parameters)^36^ and cells were embedded via UMAP. Clusters were computed using Louvain clustering (resolution 0.3). Differentially expressed (DE) genes were identified using the Wilcox Rank sum test retaining protein coding genes expressed in > 20% of cells with log2FC > 1 and adjusted p-value < 0.05. Pseudo bulk count matrices were generated per batch/cluster by averaging expression, normalized as above, and used to merge “non-informative” clusters (clusters with < 100 DE, log2FC > 1, p < 0.05, Wilcox test). GO analysis was performed based on the DE genes in R using the package ClusterProfiler (4.20.0), with ontologies queried from the “Org.Hs.Eg.db” package. Redundant terms and terms above the 0.05 q-value cut-off were excluded from consideration.

### Echocardiography

Mice were anesthetized by intraperitoneal injection with a mixture of ketamine (100 mg/kg) and xylazine (10 mg/kg). After shaving the thoracic area, the mouse was fixed in a supine position on the pre-warmed imaging platform integrated within the Vevo Animal Monitoring System (FUJIFILM VisualSonics Inc., Toronto, ON, Canada). During imaging mouse respiratory rate and temperature were closely monitored. Using the Vevo F2 ultra-high-frequency, real-time ultrasound system (FUJIFILM VisualSonics Inc., Toronto, ON, Canada) equipped with a 71-30 MHz center frequency linear array transducer (UHF71x, FUJIFILM VisualSonics Inc., Toronto, ON, Canada), high-resolution echocardiographic images were acquired under blinded conditions as previously described.^37^ Likewise, all analyses were conducted by another blinded operator using the Vevo LAB software version 5.8.2 (FUJIFILM VisualSonics Inc., Toronto, ON, Canada) by a single trained individual as previously described.^37^

### MI induction and iPSC-CM transplantation

Mice were anesthetized as described above, and re-shaved if fur had regrown in the thoracic area. After intubation and ventilation mice were fixed on a pre-warmed surgical field to perform a left-sided thoracotomy followed by MI as previously described.^38^ In brief, the pericardium was ruptured and the left anterior descending coronary artery (LAD) was ligated using an 8.0 Prolene suture. Hearts were then injected two times in the infarct border zone with 5 µL of either i) Vehicle (PBS^Cocktail^: PBS/1% hydrogel/0.1% Y27632 (2HCl)), ii) iPCS-vCM^SC^ in PBS^Cocktail^ (20,000 cells/µL), or iii) iPSC-vCM^Sphere^ in PBS^Cocktail^ (20,000 cells/µL corresponding to 80 spheres/µL). Finally, thoracic muscle layers were carefully closed with a 6.0 Prolene suture whereafter the skin was closed with a 5.0 Ethicon Vicryl suture. Post-surgery treatment with subcutaneous injections of saline to prevent dehydration and a dose of extended-release Buprenorphine Ethiqa XR (3.25 mg/kg) for analgetic treatment. At study termination, the mice were sacrificed by cervical dislocation, and the heart was excised and prepared for histology while lung, liver, spleen, and kidney were snap-frozen for AluY detection.

### Histology

Dissected hearts were immediately rinsed in PBS with heparin, then fixed overnight in 4% Neutral Buffered Formalin (NBF), rinsed in PBS, before dehydration in ethanol and paraffin embedding. For stainings, sections were deparaffinized as previously described.^37^ Before, hematoxylin and eosin staining, sections were fixed in 4% neutral buffered saline (NBF), rinsed in running tap water, stained with hematoxylin followed by rinse and development in running tap water. Finally, sections were stained with eosin, dehydrated in ethanol, and mounted with Pertex. To measure infarct size, we performed Masson’s Trichrome staining and quantification as previously described.^37,38^ Immunofluorescence staining was performed as previously described.^39^ (For primary and secondary antibodies see Table S1). Mounting was performed using DAPI (Vectashield cat.no. H-1200). Confocal imaging was performed using a Leica TCS SP8 inverted microscope (Leica Microsystems, Wetzlar, Germany), equipped with an HC PL APO 40x/1.10 water objective and PMT and HyD detectors. Images were acquired using sequential scanning in Leica Application Suite X (Las X) v3.x software, at 1,024×1,024 pixel resolution using 405, 488, 553, and 647 nm laser lines, with laser power and scan speed kept constant across all samples. Images were processed in Adobe Photoshop and adjustments (brightness/contrasts) were applied equally to all sample sections to enable visualization of all channels in the merged pictures.

### qPCR

Total RNA was isolated from collected iPSC-CMs using TriReagent as previously described,^40^ whereas cDNA synthesis and qualitative real time polymerase chain reaction (qRT-PCR) were performed as reported.^41^ QRT-PCR was run on a 7900HT Fast Real-time PCR system with the following conditions: holding for 10 min at 95°C, followed by 40 cycles consisting of 15 sec of denaturation at 94°C, 30 sec of annealing at 60°C and 30 sec of elongation at 72°C (for primer sequences and precise annealing temperature see Table S2). The data was analyzed as previously described^40^ by normalization to at least two stably expressed endogenous genes (GAPDH and PGK1) according to the qBase Plus platform (Biogazelle/CellCarta).

### AluY

Genomic DNA was isolated from the snap-frozen tissue or iPSC cells using TriReagent. Briefly, following defrosting, a suitable amount of TriReagent was added to the tissue in a M tube (Miltenyi) and tissue was dissociated using a gentleMACS dissociator as recommended by manufacturer. Polyacrylic carrier was added before addition of 1-Brom-3-Chloro-Propene. After phase separation, DNA was pelleted and rinsed (x2) in 0.1 M trisodium/10% ethanol, (x1) in 75% ethanol, and then centrifugated, before air-drying and washing in TE buffer. Collected DNA was quantified using a DeNovix DS-11 series spectrophotometer/fluorometer. For detection of the primate specific AluY sequence (primer sequences, see Table S2), TaqMan qRT-qPCR was performed according to the manufactures recommendations and was run on the 7900HT instrument under the following conditions: holding for 10 min at 95°C, followed by 40 cycles consisting of 15 sec at 95°C, and 60 sec at 60°C. For each tissue, calibration curves were generated by spiking fixed amounts of genomic iPSC DNA into gDNA from the respective tissue and using the gDNA mix as template for qPCR. The Limit of Detection (LOD) was determined to be below 0.00069% iPSCs regardless of tissue, equaling less than one human cell pr. mg of mouse gDNA.

## ACKNOWLEDGEMENTS

We would like to thank Charlotte Nielsen and Tina K. Andersen (Andersen group, Odense University Hospital/University of Southern Denmark) for their excellent technical assistance throughout this study. Confocal image acquisition was performed at the Danish Molecular Biomedical Imaging Center (DaMBIC, University of Southern Denmark), supported by the Novo Nordisk Foundation (NNF) (grant agreement number NNF18SA0032928). Figure 1A, 2D and 3C were created with BioRender.com.

Financial support for this research was received from The Novo Nordisk Foundation (#NNF21OC0071952), the Lundbeck Foundation (#R313-2019-573), Danish Cardiovascular Academy (#PD2Y-20214004), Danish research council (#8045-00019B, Sapere Aude) and (#4285-00027B), Strategic research finance (“Flagship program” #001) from Odense University Hospital and University of Southern Denmark and PhD scholarships for AKST (Region of Southern Denmark (#2025-0248), Clinical Research Institute/University of Southern Denmark, and Odense University Hospital (#7079)); SML (Danish Cardiovascular Academy (#PHD2025016-DCA) and FAB (Innovation Foundation Denmark (#2052-00021B)).

## AUTHOR CONTRIBUTIONS

AKST, DGE: Collection of data, data analysis and interpretation, manuscript writing, and final approval of manuscript; SBM, SML, FAB, BGA, FGW: Collection of data, data analysis and interpretation, and final approval of manuscript; CHJ: Data analysis and interpretation, manuscript editing and final approval of manuscript; DCA: Conception and design, collection of data, data analysis and interpretation, manuscript writing, final approval of manuscript, and financial support.

## DECLARARION OF INTEREST STATEMENT

The authors declare no conflicts of interest.

