## Supplemental material for "Poor survival after myocardial infarction in immune deficient NXG B2m mice is alleviated by transplantation of induced pluripotent stem cell derived cardiomyocytes (iPSC-CM) formulated as spheres, but not single cells"

A

| Metric | Batch1 | Batch2 | Batch3 | Batch4 |
|---|---|---|---|---|
| # Cell | 14,258 | 10,000 | 12,413 | 11,362 |
| # Total reads | 230,138,635 | 507,950,057 | 373,054,948 | 325,837,084 |
| Median UMI counts per cell | 1076 | 1735 | 2282 | 6112 |
| Fraction of reads in cell | 0.74 | 0.79 | 0.74 | 0.73 |
| Q30 bases in read | 87.9 | 91.9 | 91.6 | 93.1 |
| Reads mapped confidently to the transcriptome | 41.9 | 55.4 | 49.2 | 54.5 |

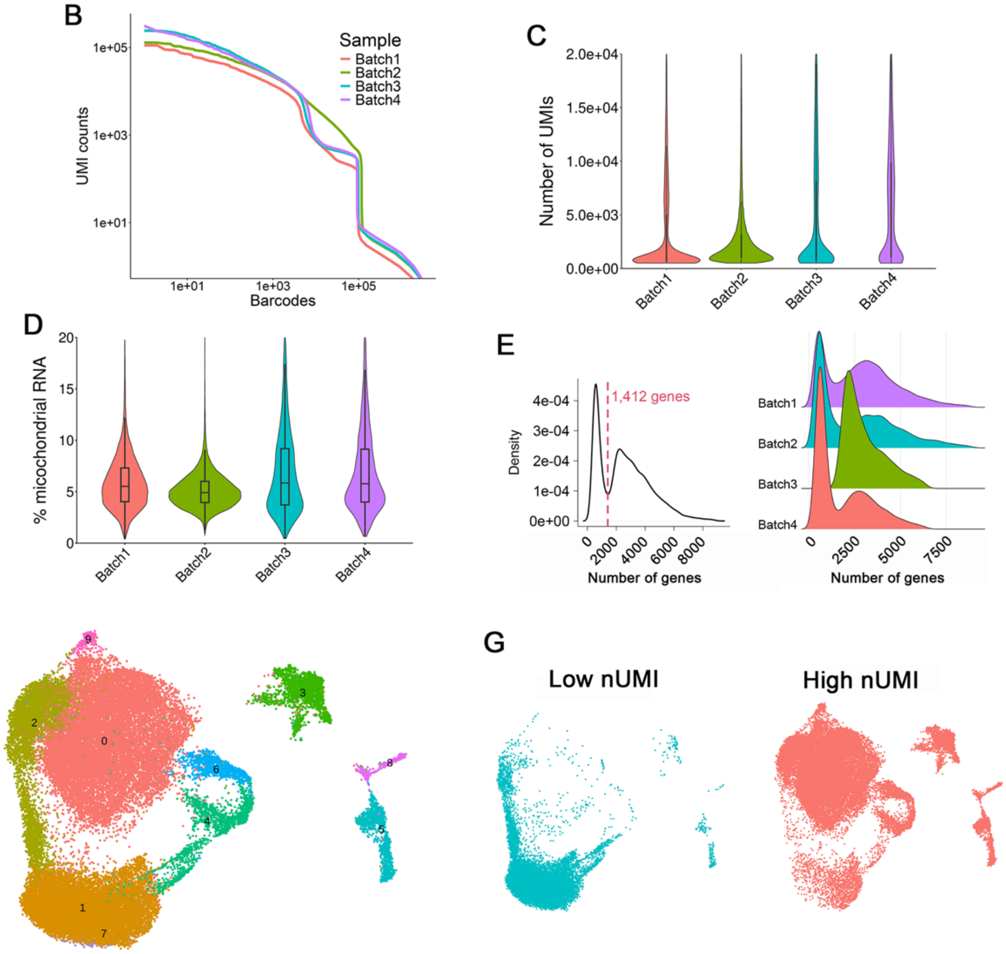


**Figure S1. Quality control of the four samples analyzed by scRNA-seq**
(A) # Cells, # Total reads, Median Unique molecular identifier (UMI) counts per cell, Fraction of reads in cell, Q30 bases in read, and Reads mapped confidently to the transcriptome for the four batches. (B-D) Quality analyses of the four batches. (B) Barcode rank plot (knee-plot) showing log scaled barcodes ranked by number of UMI vs log scaled UMI counts. (C) Median UMI counts per cell. (D) Percentages of mitochondrial transcripts. (E) Density plot showing the distribution of the number of genes per cell in the collected dataset (left) and separated by batch (right) (F) uniform manifold approximation and projection (UMAP) embedding showing cells prior to filtering based on UMI count, colored by Louvain clusters and (G) classified by the number of UMIs group.

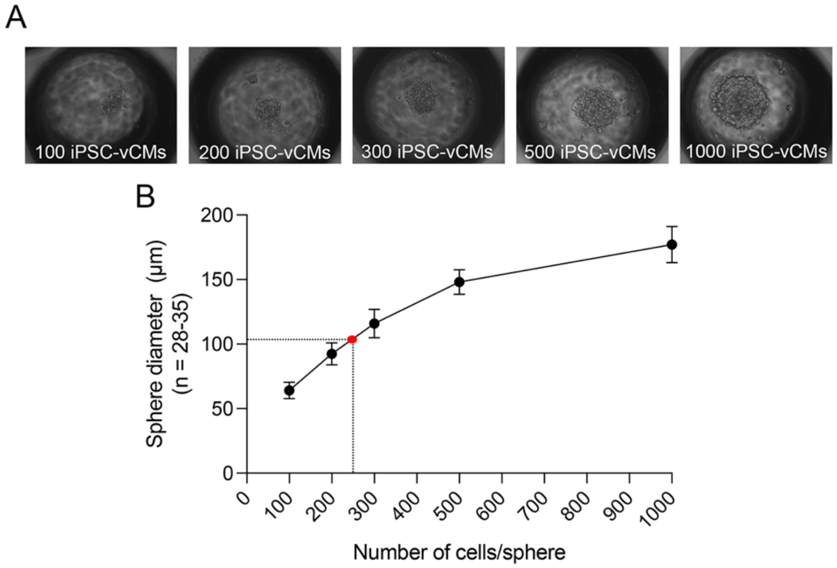


**Figure S2. Optimal numbers of iPSC-vCMs for obtaining iPSC-vCM$^{Spheres}$ with a diameter suitable for transplantation**

(A) Representative pictures of spheres consisting of 100, 200, 300, 500, or 1000 iPSC-vCMs, respectively. (B) Sphere diameter of spheres consisting of 100, 200, 300, 500, and 1000 iPSC-vCMs, the red dot and the dashed lines indicate that 250 iPSC-vCMs are necessary to form spheres with an approximately diameter of 100 µm.

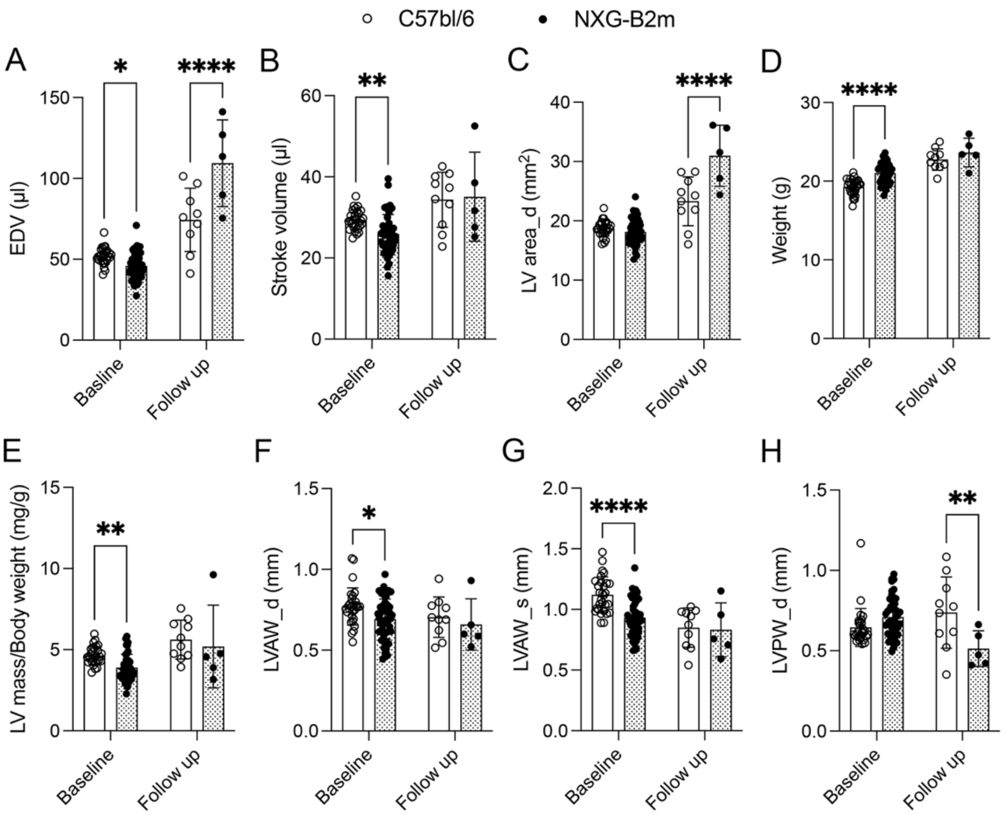


**Figure S3. Heart function in C57bl/6 and NXG B2m mice at baseline and following myocardial infarction (MI)**
(A) End diastolic volume (EDV), (B) Stroke volume, (C) Left ventricle (LV) area in diastole, (D), weight, (E) LV mass/Body weight, (F) left ventricle anterior wall (LVAW) thickness in diastole, (G) LVAW thickness in systole, and (H) left ventricular posterior wall (LVPW) thickness in diastole in C57bl/6 and NXG-B2m mice at baseline and follow up after myocardial infarct (MI). *≤0.05: **: p<0.01, ****: p<0.0001.

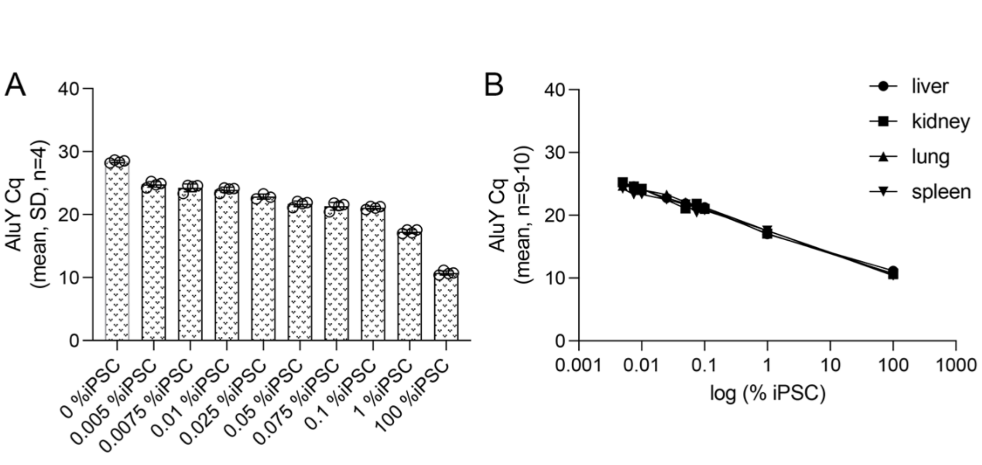


**Figure S4. Sensibility of the AluY qPCR biodistribution assay**

(A) AluY qPCR Cq-values of gDNA samples (from mouse liver, -kidney, -lung and -spleen) containing from 0 to 100 percentage of human induced pluripotent stem cells (iPSC) gDNA. (B) AluY qPCR Cq-values from the mixed gDNA samples with mouse liver, kidney, lung, and spleen each depicted as a function of log (% iPSC) and showing a linear relationship. (C) AluY Cq-values of samples containing 0.00069%, and 0.005% iPSC gDNA showing significant differences to the 0% control sample.

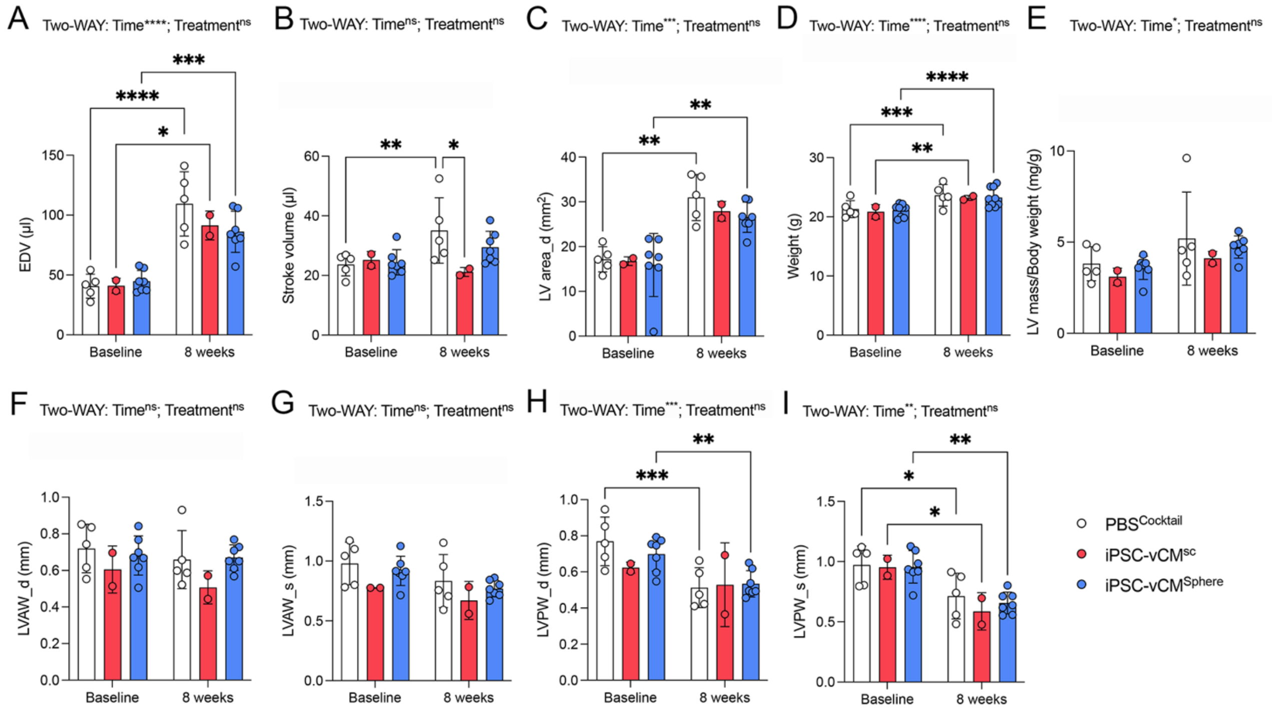


**Figure S5. Heart function of NXG B2m mice at baseline and following myocardial infarction (MI)**

(A) End diastolic volume (EDV), (B) stroke volume, (C) left ventricular (LV) area in diastole, (D) weight, (E) LV mass/Body weight (F) left ventricular anterior wall (LVAW) thickness in diastole, (G) LVAW thickness in systole, (H) left ventricular posterior wall (LVPW) thickness in diastole, and (I) LVPW thickness in systole in NXG B2m mice at baseline and 8 weeks after myocardial infarct and injection of $PBS^{Cocktail}$, $iPSC\text{-}vCM^{SC}$ or $iPSC\text{-}vCM^{Sphere}$. *≤0.05: **: p<0.01, ***: p<0.001, ****: p<0.0001.

**Table S1. List of primary and secondary antibodies used.**

| | Catalogue number | Company | Dilution |
|---|---|---|---|
| **Primary antibodies** | | | |
| Rabbit anti-Ku80 | 2180 | Cell Signalling Technology | 1:200 |
| Mouse anti-Myosin heavy chain Type 1 (MYH7) | BA-D5 | DSHB | 1:200 |
| Mouse anti-CX43 | SAB4200730 | Sigma-Aldrich | 1:250 |
| Mouse anti-$\alpha$-actinin | A7811 | Sigma-Aldrich | 1:200 |
| Rabbit anti-GFP | Ab290 | Abcam | 1:500 |
| Mouse anti-tropomyosin | T9283 | Sigma-Aldrich | 1:500 |
| Rabbit anti-Nkx2.5 | Sc-14033 | Santa Cruz | 1:100 |
| Sheep anti-Mef2c | AF6786 | BioTechne | 1:40 |
| **Secondary antibodies** | | | |
| Donkey anti-mouse IgG 555 | A-31570 | Invitrogen | 1:200 |
| Donkey anti-rabbit IgG 488 | A-21206 | Invitrogen | 1:200 |
| Donkey anti-sheep IgG 488 | A-11015 | Invitrogen | 1:200 |
| Donkey anti-rabbit IgG 647 | A-31573 | Invitrogen | 1:200 |

**Table S2. Supplemental Table 2. Human specific qPCR primers used.**

| Name | Sequence | Sample volume (ng) | Annealing temp (°C) |
|---|---|---|---|
| *GAPDH* | F: GCCACATCGCTCAGACACCATGG<br>R: TCCCGTTCTCAGCCTTGACGGT | 2 | 60 |
| *PGK1* | F: GTCGGCTCCCTCGTTGACCGAA<br>R: GGGACAGCAGCCTTAATCCTCTGGT | 2 | 60 |
| *OCT4* | F: CAGTGCCCGAAACCCACAC<br>R: GGAGACCCAGCAGCCTCAAA | 2 | 60 |
| *Sox2* | F: CCCT GTGGTTACCTTTTCCT<br>R: AGTGCTGGGACATGTGAAGT | 2 | 60 |
| *TNNT2* | F: GACAGAGCGGAAAAGTGGGA<br>R: GCGGGTCTTGGAGACTTTCT | 2 | 60 |
| *MYL2* | F: TGTCCCTACCTTGTCTGTTAGCCA<br>R: ATTGGAACATGGCCTCTGGATGGA | 2 | 60 |
| *MYL7* | F: ACATCATCACCCATGGAGACGAGA<br>R: GCAACAGAGTTTATTGAGGTGCCC | 2 | 60 |
| *FABP3* | F: CACTCGCACTTATGAGAAAGA<br>R: AGGAAGAAATGAGGCAATGT | 2 | 60 |
| *SCN5A* | F: TCTTCACAGGCGAGTGTATTG<br>R: GACAACCACGAAGTCGAAGATA | 2 | 60 |
| *CKMT2* | F: GGAGAGAGGCCAAGATATTAAG<br>R: CGTACCAGCAGACAGATTATT | 2 | 60 |
| *SiRPA* | F: ACCTGGCTCAGGCTAGTTCCAAAT<br>R: TGTGCACACGTATGTGCTGTCTCT | 2 | 60 |
| *P53* | F: AGCACTGTCCAACAACACCA<br>R: CTTCAGGTGGCTGGAGTGAG | 2 | 60 |
| *BAX* | F: CCCTTTTGCTTCAGGGTTTCAT<br>R: GGAAAAAGACCTCTCGGGGG | 2 | 60 |
